# Drug target discovery via network modeling: a mathematical model of the *E. coli* folate network response to trimethoprim

**DOI:** 10.1101/712257

**Authors:** Inderpreet Jalli, Sophia Lunt, Wenjia Xu, Carmen Lopez, Andreas Contreras, Cari-Sue Wilmot, Timothy Shih, Frederik Nijhout

**Affiliations:** Department of Biology, Duke University, Durham, North Carolina, United States of America; Department of Biology, Michigan State University, East Lansing, Michigan, United States of America; Department of Earth and Atmospheric Science, University of Houston, Houston, Texas, United States of America

**Author notes:** Corresponding author (IJ). These members contributed equally to this work (Inderpreet Jalli, Sophia Lunt, Frederik Nijhout). These members also contributed equally to this work (Wenjia Xu, Carmen Lopez, Andreas Contreras, Cari-Sue Wilmot, Timothy Shih).

## Abstract

The antibiotic trimethoprim targets the bacterial dihydrofolate reductase enzyme and subsequently affects the entire folate network. We present an expanded mathematical model of trimethoprim’s action on the *Escherichia coli* folate network that greatly improves upon Kwon *et al.* (2008). The improvement upon the Kwon Model lends greater insight into the effects of trimethoprim at higher resolution and accuracy. More importantly, the presented mathematical model enables drug target discovery in a way the earlier model could not. Using the improved mathematical model as a scaffold, we use parameter optimization to search for new drug targets that replicate the effect of trimethoprim. We present the model and model-scaffold strategy as an efficient route for drug target discovery.

## Introduction

### Antibiotic resistance

Antibiotic resistance is a major health policy concern. New and resistant forms of common infections such as tuberculosis necessitate urgent drug development efforts [1–3]. Strategies such as discovering new bacterial communication networks and inhibiting these networks are a popular avenue for drug discovery yet are very expensive in both time and resources. Utilizing math models to gain a greater level of insight into the mechanisms of known drugs may offer opportunities for drug development, using well-studied and richly described pathways that already show weakness to chemical intervention to search for new potential targets [4]. The presented work models the mechanism of a common antibiotic, trimethoprim, at the biological information layer at which it functions, the metabolic network level. This is done not only to study the mechanism of trimethoprim but to find alternatives to it by attempting to replicate its effect on the folate network, which is known to be critical to cell function. The presented mathematical model improves on previous work by providing a higher resolution model of Trimethoprim’s effect on *E. coli*. Without a highly detailed model of the bacterial folate network and trimethoprim’s effect on it, the presented strategy of drug target discovery would not be possible.

### The folate network and trimethoprim

The folate network is a traditional therapeutic target for both cancerous and bacterial cells due to the integral role folates play in cell division [5, 6]. The folate network provides and accepts one-carbon units for the biosynthesis of amino acids and metabolites such as S-adenosyl methionine (SAM), the universal methyl group donor [7–9]. The antibiotic trimethoprim (TM) inhibits the activity of bacterial dihydrofolate reductase (DHFR), an enzyme that converts dihydrofolate (DHF) to tetrahydrofolate (THF, Fig 1). DHFR inhibition causes a spike in DHF. DHF in turn inhibits folypolyglutamate gamma synthetase (FPGS), the enzyme tasked with adding glutamates to THF and its derivatives [10, 11]. Because folate-catalyzed conversions of one-carbon units are sensitive to glutamation levels of folates, the inhibition of FPGS disrupts critical cell functions [11–14].

**Fig 1.**
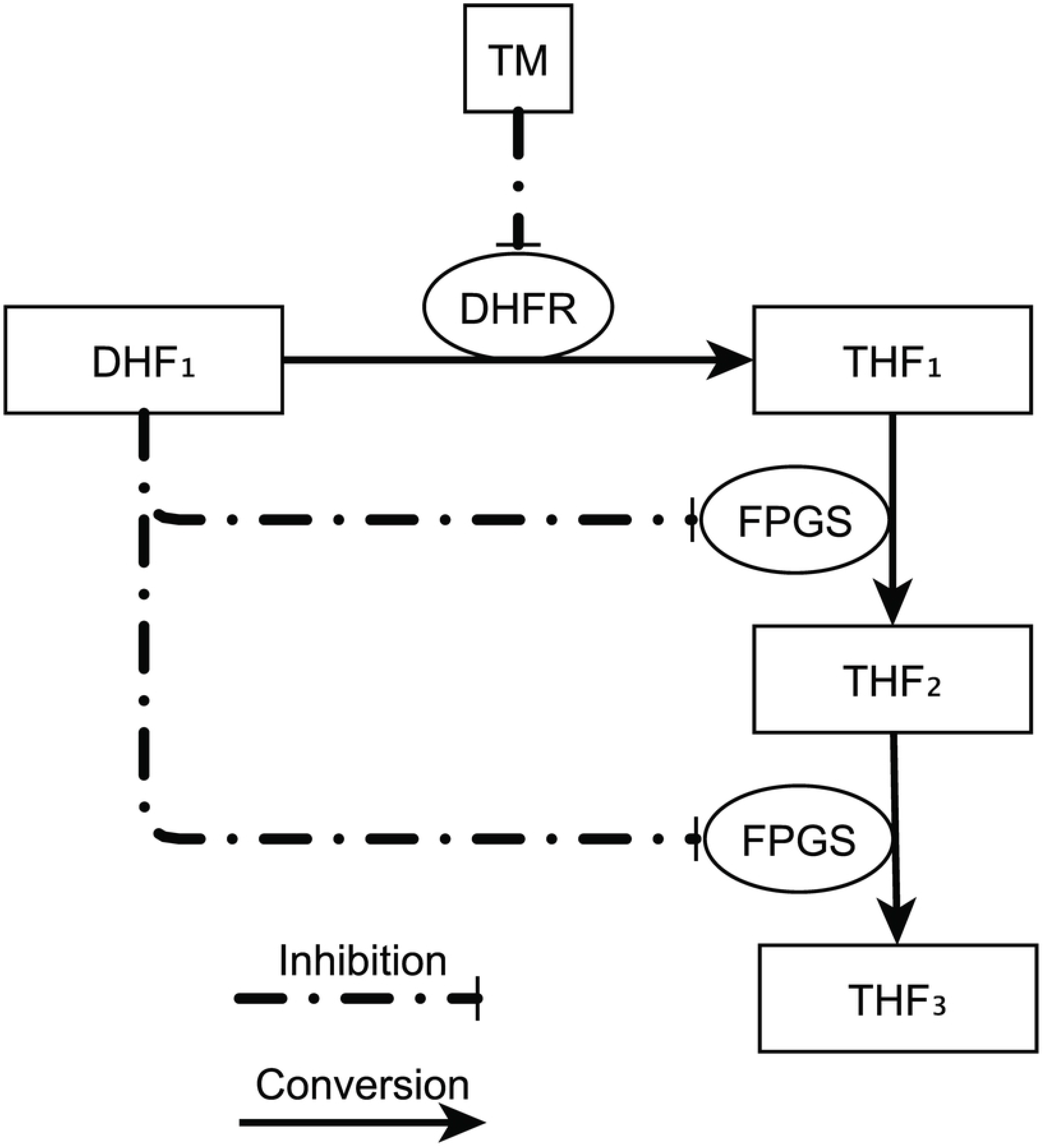
A simplistic diagram of effect of trimethoprim on DHFR and FPGS, roughly approximated in Kwon Model. Boxes = metabolites or antibiotics. Ovals = enzymes, connected to solid lines with arrows indicating metabolite conversions. Dashed lines = inactivation. Trimethoprim inhibits DHFR, which converts DHF_1_ to THF_1_, leading to a spike in DHF_1_. FPGS adds a glutamate to THF_1_, and THF_2_, converting each to THF_2_, and THF_3_, respectively.

### Folate network models and drug target search

Kwon *et al.* wrote a mathematical model of the disruptive effect of TM focused on folate interconversions which roughly has the structure and resolution of Fig 1, and which we refer to as the *Kwon Model* [10]. The *Kwon Model* compressed all derivatives of THF into three variables, but Kwon et al. recorded experimental data for many THF derivatives. In this work we present a higher resolution version of the *Kwon Model* referred to as the *TM Model*, that exploits the experimental data recorded by Kwon et al. The structure of the *TM Model* is shown in Fig 2. We then use the current *TM Model* as a scaffold for drug target searching by replicating the effects of TM on the *E. coli* folate network without inhibiting DHFR or FPGS. In doing this we create a *Rewired Model* that shows a simulated time-course progression similar to the *TM Model*, but without TM in the simulation. This overall strategy is outlined in Fig 3. The *TM Model* by the nature of its higher resolution, higher number of enzyme and metabolite nodes, and higher accuracy of describing the effect of TM on the folate network, allows for TM alternatives to be explored in a way that the *Kwon Model* cannot. Since multiple interconversions of THF_n_ and DHF_n_ are described in the *TM Model*, substituting TM with other possible disruptors of the folate network is feasible, guided by Kwon et al.’s experimental data. This approach allows us to propose new targets for antibiotic activity that would be advantageous against TM-resistant bacteria, and provides a guide for future work on other systems that evolve in response to chemical therapeutics [15, 16].

**Fig 2.**
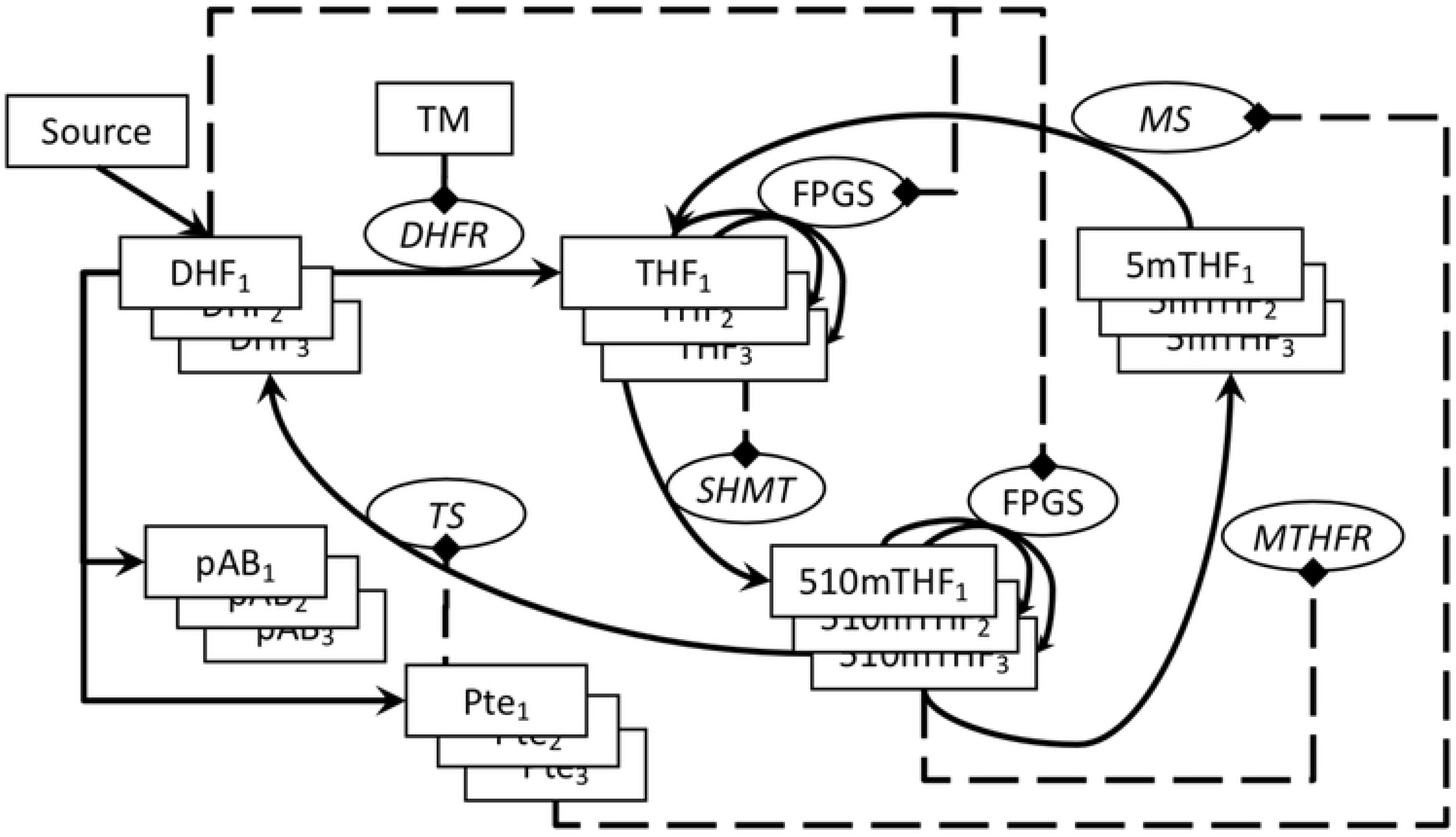
TM Model of E. coli folate network. (Ovals = enzymes modeled with Michaelis-Menten equations; boxes = metabolites, TM, or source; dashed lines = inhibitions; straight lines without attached oval = linear conversions. The source (top left) represents 7,8-dihydropteroate, and mass input is modeled as a linear conversion into DHF_1_ only by dihydrofolate synthase (DFHS). Once DHF_1_ is produced, it is converted into THF_1_ via DHFR. TM inhibits all DHFR activity. THF_1_ is converted into THF_2_ and then to THF_3_ by FPGS, which also works on all THF_n_ derivatives (except 5MTHF_n_), indicated by thin solid lines. Only FPGS carries out interconversions between glutamation states. All FPGS activities have independent parameter sets. DHF_n_ inhibits FPGS activity. TS converts 510MTHF_n_ to DHF_n_. SHMT converts THF_n_ to 510MTHF_n_, and MTHFR converts 510mTHF_n_ to 5MTHF_n_. MS does not convert monoglutamates of 5MTHF_n_ to THF_n_, but its arrow is shown as other enzymes for simplicity. DHF_n_ is converted into pAB_n_ and Pte_n_, which act as sinks for the system. Experimental concentration data exists for all metabolites shown in the figure. Pte_n_ inhibits TS and MS, THF_n_ inhibits SHMT, and 510MTHF_n_ inhibits MTHFR. These inhibitions however are not as strong as that of TM on DHFR. Formyl THF_n_ derivatives were not included due to a lack of experimental data.

**Fig 3.**
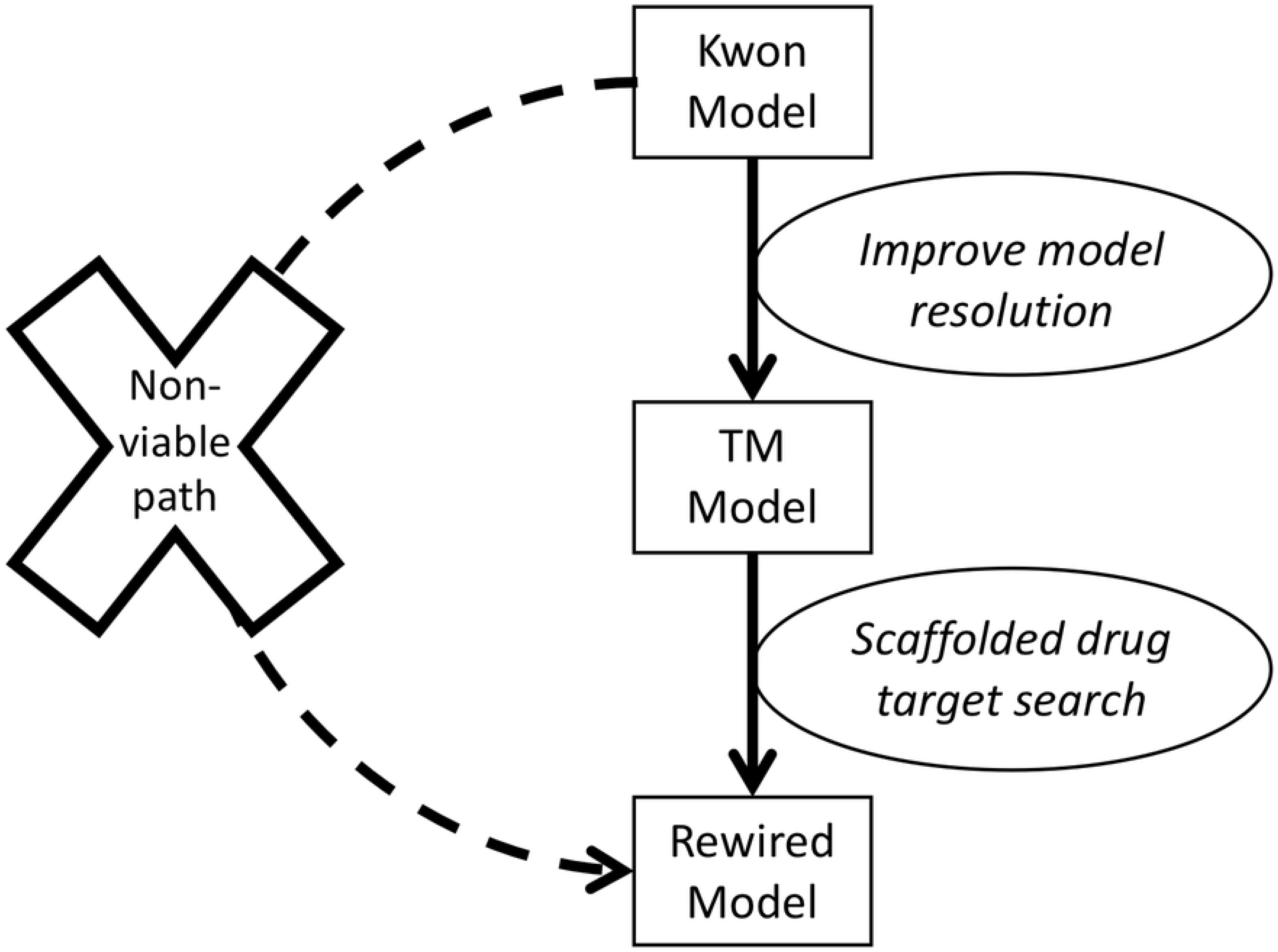
Drug target searching requires high resolution models. The Kwon Model is improved upon to create the TM model, which has more metabolite and enzyme nodes. The TM model describes the effects of TM at high resolution and accuracy. The TM model, once developed, is used as a scaffold for drug target searching (Rewired Model). The Kwon Model, due to having only two enzymes (approximated in Fig 1) and compressing all forms of THF into just three nodes, is not viable for drug target discovery (pathway on left).

### Abbreviations

Enzymes are listed with abbreviation, name, and Uniprot ID and/or EC number. A subscript indicates the number of glutamates added onto a molecule. Terms such as DHF_n_ refer to molecules that vary only in number of glutamates. DHFR: dihydrofolate reductase (P0AFS3 / EC: 1.5.1.3), DHFS: dihydrofolate synthase (P08192 / EC: 6.3.2.12), SHMT: serine hydroxymethyltransferase P0A825/ 2.1.2.1), MTHFR: methylenetetrahydrofolate reductase (P42898 / 1.5.1.20), TS: thymidylate synthase (P0A884 / EC: 2.1.1.45), METH: methionine synthase (P13009 / EC: 2.1.1.13), METE/MS: B12 independent methionine synthase (P25665 / EC: 2.1.1.14), FPGS: folypolyglutamate synthetase (P08192 / EC: 6.3.2.17), MET: L-methionine, SAM: S-adenosyl-L-methionine, SRH: S-ribosyl-L-homocysteine, SAH: S-adenosyl-L-homocysteine, HCY: L-homocysteine, TM: trimethoprim, Pte_n_: folate glutamate, pABA: para-aminobenzoate, pAB_n_: para-aminobenzoylglutamate, DHF_n_: dihydrofolate, DHP: dihydropteroic acid, THF_n_: tetrahydrofolate, 5MTHF_n_: 5-methyl-THF, 510MTHF_n_: 5,10-methylene-THF. Enzyme parameters are referred to with enzyme abbreviation followed by the type of parameter, then the glutamate that is being acted upon, e.g. DHFRKM1 refers to DHFR’s *Km* constant for DHF_1_. The included models are heavily focused on the differing activities of enzymes on a variety of unique glutamations of molecules such as THF_n_. The velocities of enzymes are denoted by the enzyme abbreviation and then the glutamation level of the general molecule they are named for acting upon. For example, DHFR1 refers to the velocity of DHFR’s activity on DHF_1_. In the case of FPGS which works on THF_n_ and its derivatives with differing affinities (except for 5MTHF_n_), the substrate and glutamate are used in the velocity or parameter abbreviation: FPGSTHF1 refers to FPGS activity on the THF_1_ substrate. SSE refers to sum of squared errors, a common metric for describing the accuracy of a prediction to the real world data.

## Materials and Methods

### Experiments

Folate concentrations were measured absolutely as described in Kwon *et al*. [10, 11]. Time course progression of folate concentrations and their glutamations can be seen in Fig 4. All rates and velocities are shown in μM/minute. TM was added to *E. coli* growth medium at O.D. ~0.5 at a concentration of 4 μg/mL, immediately after time zero data point collection [10]. *E. coli* strain NCM3722 was used for all experiments [10]. Cells were grown in Gutnick minimal salts medium (Sigma-Aldrich), in a shaking flask at 37°C.

**Fig 4.**
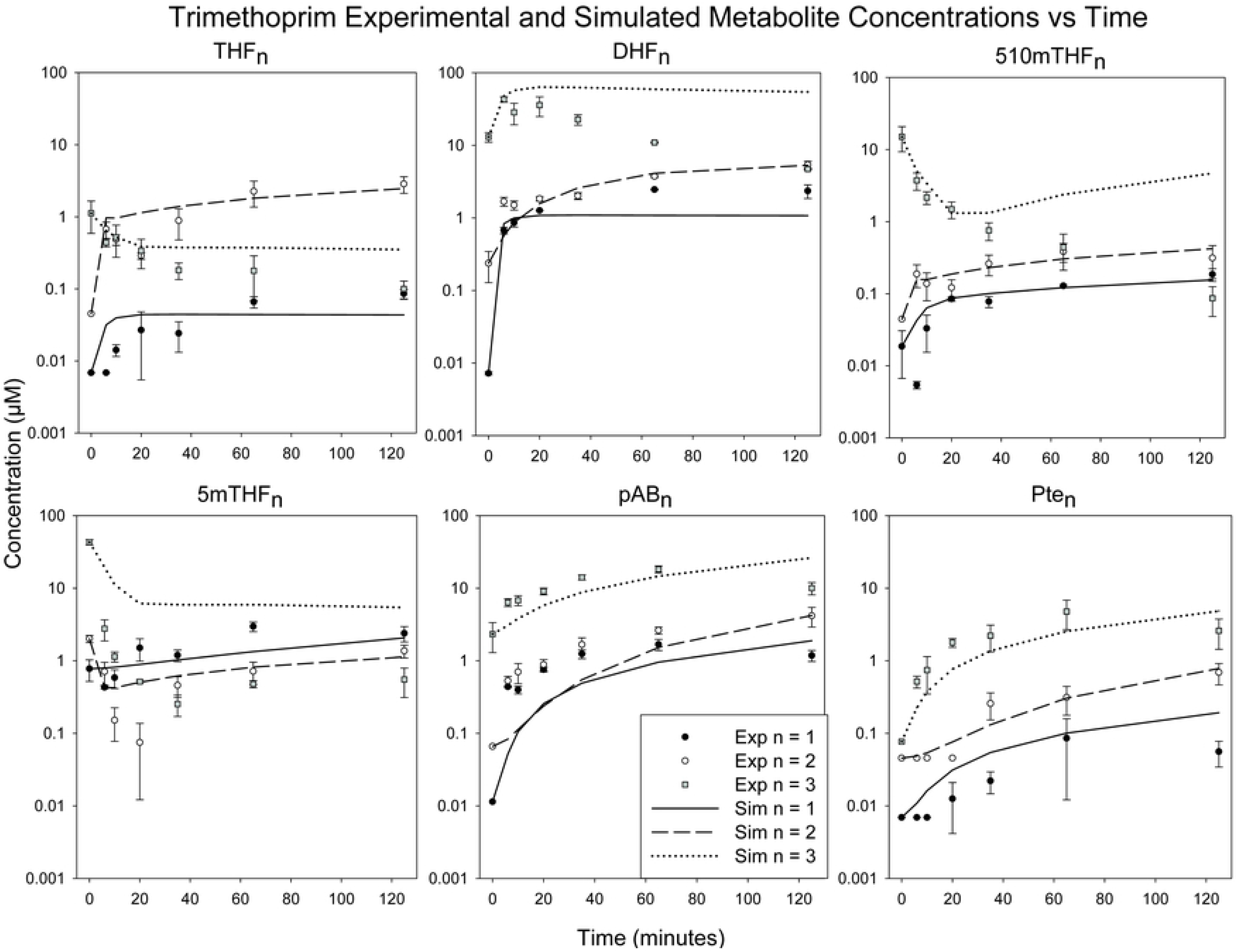
Folate disruption and simulation of the TM Model. TM in growth media causes a severe deviation from initial folate concentrations (dots with error bars). THF_n_ progression shows high similarity to experimental progression (SSE = 1.83, with THF_2_ SSE = 1.64). DHF_3_ similarity is poor (SSE = 8063), creating bulk of error for model (SSE total = 8947). The experimental time course metabolite concentration data can be found in the data in S3 Table.

### Mathematical modeling of the *TM Model*

All simulations and calculations were carried out in Matlab version 2014a (Simbiology Toolbox), on an Intel Q8200 2.33GHz processor, using Microsoft Windows 7, 64-bit, and ODE solver ode15s. Fig 2 displays the structure of the *TM Model*. Variables are concentrations of DHF_n_, pAB_n_, Pte_n_, THF_n_, and all THF_n_ derivatives, at three glutamation levels. Variable time-course progression is fitted to experimental data shown in Fig 4. The *Kwon Model* summated all THF_n_ and THF_n_ derivatives into three variables: THF_1_, THF_2_, and THF_3_. The *Kwon Model* had a lower number of enzyme kinetics equations describing interconversions of DHF_n_ and THF_n_ [10]. In the current *TM Model*, THF_n_ and THF_n_ derivatives are treated as independent variables, requiring a higher number of detailed enzyme kinetics equations to describe interconversions. This treatment of THF_n_ derivatives as unique variables allows for high resolution modeling and the creation of the *Rewired Model*. Creating the *Rewired Model* would not be possible at the resolution of the *Kwon Model* because insufficient enzyme nodes exist for a parameter search. Conversion of DHF_n_ to THF_n_ by DHFR was modeled by a variant of the Michaelis-Menten equation featuring competitive inhibition:

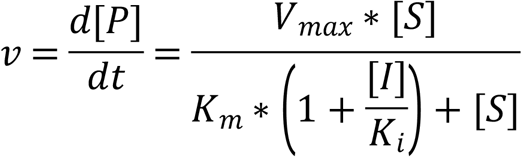

Where *Km* is the Michaelis constant, [S] is the concentration of the substrate, *K*_*i*_ is the inhibition constant for TM, [I] is the constant concentration of TM, and *Vmax* is the maximum rate achieved by the enzyme. TM was added to simulations at 40 seconds to reflect experimental protocol. Kwon *et al*. measured a significant reduction of input flux after addition of TM, which is included in the model unless otherwise stated [10]. If the Michaelis-Menten equation or a variant thereof was not used for a metabolite conversion, a linear rate equation was employed using the form presented below. In the equation below, *Ks* represents a linear transform of the substrate into a product:

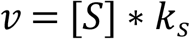

### Parameters and parameter estimation in the *TM Model*

Parameters are either Michaelis-Menten constants (*Km, Ki, Vmax*) or linear conversion constants (*Ks*). Some parameters are estimated due to insufficient experimental data in published literature, or estimated from an initial data set taken from literature, such as IC_50_ (half maximal inhibitory concentration) values. Estimated parameters were determined by fitting simulations to experimental data via sequential application of the Matlab genetic algorithm and fmincon functions.

### Creating the *Rewired Model* to explore alternatives to TM

To find an alternative to TM, we first removed the presence of TM in the *TM Model*. The resultant model with all further modifications is referred to as the *Rewired Model*. We then inserted a set of *in silico* enzyme inhibitors into the *Rewired Model* via parameter modification of existing Michaelis-Menten equations with the aim of creating a simulated time-course folate concentration progression of the same nature to that caused by TM. The experimental data used to fit the *Rewired Model* was the same as that used for the *TM Model*. DHFR and FPGS were not targeted with *in silico* manipulation in the *Rewired Model* in order to find a true alternative for TM-resistant bacteria. Parameter optimization functions were used to find the proper combination of *in silico* inhibitor. Inhibitors were simulated by introducing alterations of the *Km* and *Vmax* parameters used in the *TM Model*.

## Results

### Trimethoprim affects polyglutamation – *TM Model*

TM, which is added at 40 seconds into the model simulation, dramatically alters folate pools in the *TM Model* (Fig 4) which replicates the experimental data in *Kwon et al.* with a total SSE of 8948. DHF_3_ alone has an SSE of 8063 and is responsible for most of the error. The *TM Model* greatly improves upon the simulation resolution of the *Kwon Model* featured in the same effort [10]. DHF_1_ and DHF_2_ experience large increases both experimentally and in the *TM Model*. Simulated DHF_3_ experiences a minor spike and then slowly stabilizes to near its original concentration instead of showing a drop from initial concentration. A key experimental effect of TM is seen in the model: THF_1_ and THF_2_ increase after initial drops while THF_3_ drops continuously. THF_n_ derivatives 510mTHF_n_ and 5mTHF_n_ follow similar progressions as seen in experimental data, with the exception of 5mTHF_3_. Fig 5 highlights critical reaction velocities of the *TM Model* to show the drivers of folate concentration progression. Velocity of DHFR1 shows a drop and then a slight rebound, as would be expected due to TM inhibition of DHFR activity. FPGSTHF1 velocity experiences a sudden drop due to an increase in DHF_n_ which inhibit the activity of FPGSTHF1. Parameter estimates, drawn from both experimental work and estimation *in silico*, are shown in S1 Table.

**Fig 5.**
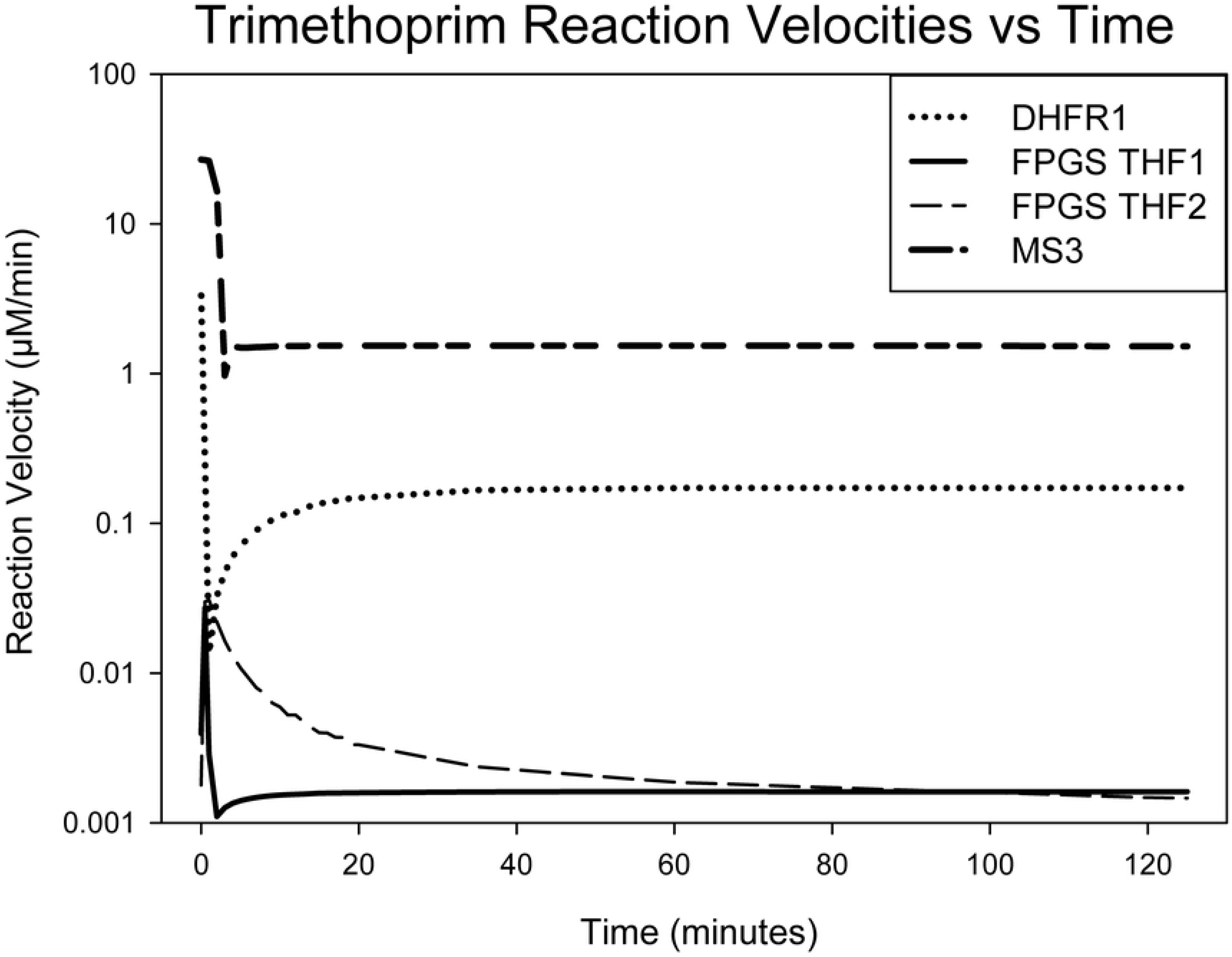
Velocities of DHFR, MS on 5MTHF_3_, FPGS on THF_1_ and THF_2_, under TM. As expected, velocities drop upon the addition of TM. Some activities such as DHFR on DHF_1_ recover slightly over time. Velocity simulations provide reference that assures drivers of simulation mirror biological processes.

### Network rewiring to explore TM alternatives – *Rewired Model*

To explore an alternative therapeutic approach to TM, a small number of enzyme inhibitors were added *in silico* while TM was removed, creating the *Rewired Model*. The *Rewired Model*’s parameters were fitted to the same experimental data used to create the *TM Model*. The simulation results of the *Rewired Model* are seen in Fig 6, and the simulated inhibitors are shown in Table 1. The total SSE of the *Rewired Model* as compared to experimental data is 46794, driven mainly by 5mTHF_3_ (SSE = 39929). All THF_n_ respond as they did in the *TM Model* although not as well *(Rewired Model SSE = 16.41, TM Model SSE = 1.83)*. DHF_n_ do not follow the *TM Model* pattern, dropping in concentration over the course of the simulation. The proposed inhibitors achieve a partial effect of TM (spiking THF_1_ / THF_2_, dropping THF_3_, alterations of 510mTHF_n_, 5mTHF_n_) without utilizing the DHF spike (further discussion below). *Rewired Model* reaction velocities are shown in S1 Fig. Most velocities except DHFR1 reach steady state early in the *Rewired Model*. In addition, velocities such as FPGSTHF1 increase instead of decreasing as was seen in the *TM Model* (Fig 5).

**Fig 6.**
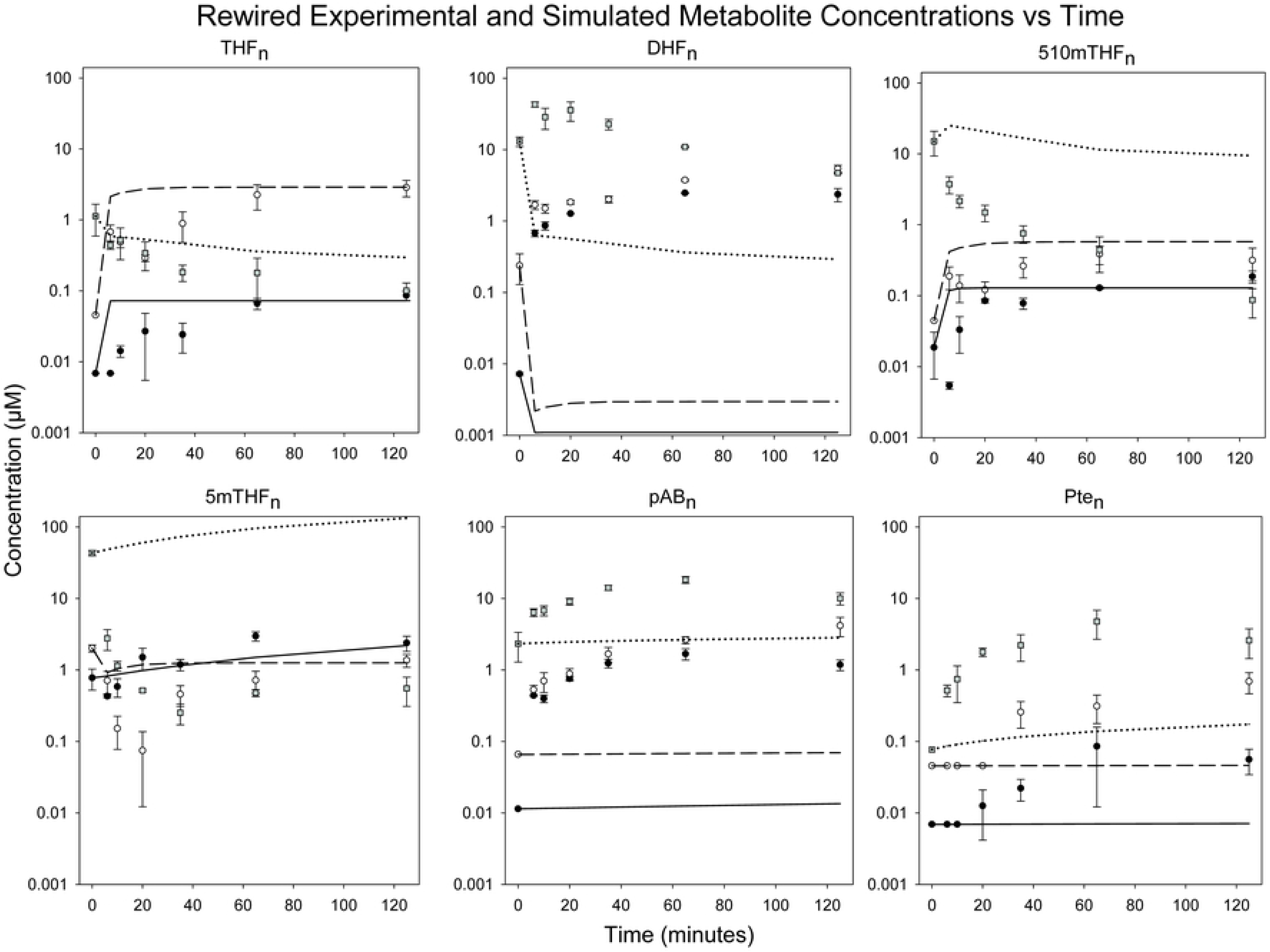
Rewired Model of E. coli folate to search for new drug targets. Simulations show an initial disruption of folate concentrations, and then a gradual stabilization. Combined THF_n_ progression shows high similarity to experimental progression.

**Table 1.** Inhibitors added in substitution of TM in Rewired Model.

| Parameters | *TM Model* | *Rewired Model* | Inhibition Type |
| --- | --- | --- | --- |
| **MSVM3** | 1.40E-01 | 1.00E-02 | Noncompetitive |
| **MTHFRKM2** | 2.99E+00 | 3.50E+00 | Competitive |
| **MTHFRKM3** | 1.11E-01 | 1.47E+01 | Competitive |
| **SHMTKM3** | 2.58E+01 | 6.80E+00 | Uncompetitive |
| **SHMTVM3** | 4.07E+00 | 3.10E+00 | Uncompetitive |

Enzymes MS, MTHFR, and SHMT are targeted with noncompetitive, competitive, and uncompetitive inhibitors respectively. TM Model parameters are shown along with inhibition type required to partially recreate effect of TM (spiking THF<sub>1</sub> / THF<sub>2</sub>, dropping THF<sub>3</sub>, alterations of 510mTHF<sub>n</sub>, 5mTHF<sub>n</sub>) in Rewired Model.

## Discussion

### The effect of trimethoprim – *TM Model*

The *TM* Model shows a more accurate and expanded view of the folate network than the previous effort in Kwon *et al*. [10]. SSE scores cannot truly be compared from *TM Model* to *Kwon Model* due to the large increase in metabolites modeled. The explicit addition of TM and THF_n_ derivatives are the key differences between the *TM Model* and the *Kwon Model*. THF_n_ derivatives 510mTHF_n_ and 5mTHF_n_ were collapsed into THF_n_ in the *Kwon Model*. In the *TM Model* they are modeled as unique variables as they exist within the bacterial cell and as the experimental data measured them. THF_n_ and its derivatives are critical to cell functionality due to their connection to the rest of cellular metabolism [8, 9].

The discrete simulation of THF_n_ and its derivatives allows for accurate modeling of THF_n_ interconversions to and from its derivative forms. These interconversions feature inhibitory interactions within the network such as Pte_n_ inhibition of TS. In addition, the inhibition of FPGS by DHF_n_ can be included, affecting FPGS’ activity on THF_n_ and all its derivatives in varying degrees (Fig 2).

An imbalance of glutamation levels across folates from the zero time point is clearly reflected in the simulation, as it is in the experimental results. The glutamation disruption of THF_n_ and its derivatives is a critical feature of the *TM Model*. TM operates by disrupting glutamation balance of folates. The spike in DHF_n_, shown in the *TM Model*, results in a sudden drop in the velocity of all FPGS activities as seen in Fig 4 and as expected by experiments on FPGS and DHF_n_ *in vitro* (Kwon et al.). FPGSTHF2 velocity shows a more relaxed response to the DHF_n_ spike, suggesting some buffering ability in the system.

The poor replication of DHF_3_ progression and triple glutamated folates overall is a weakness in the *TM Model*. We suspect it is due to experimental data on enzyme function at triple glutamation being sparse as compared to experimental data on singly glutamated folates. Better descriptions of enzyme function at this glutamation level may allow for better modeling in the future. Most importantly, the increased accuracy of the *TM Model* makes it an excellent platform for drug design.

### Network rewiring to explore TM alternatives

The effort to find a TM alternative centered on established drug design principles of using multiple antibiotic agents simultaneously and parameter optimization [17–19]. The simulated inhibitors found via optimization in the *Rewired Model* target MS, MTHFR, and SHMT instead of DHFR alone as seen in Table 1. The overall SSE is driven mainly by inaccuracy in 5mTHF_3_.

THF_n_ follows the progression seen in the *TM Model*, and DHF_n_ drops in concentration in the *Rewired Model* instead of increasing as it does in the *TM Model*, which is potentially problematic, since DHF_n_ is a primary driver of the effect of *TM*. However, our objective is to find an alternative and get a disruption in glutamation mix of folates that disrupt cellular metabolism. Our goal is not simply to replicate the domino effect of TM, which causes a spike in DHF_n_ and then inhibits FPGS. 510mTHF_n_ and 5mTHF_n_ generally (with exception of 5mTHF_3_) follow the same time course progression in the *Rewired Model* as in the *TM Model*, which is encouraging. THF_n_ and its derivatives are more critical to cell functionality due to their connection to the rest of cellular metabolism [8, 9]. The drop in DHF_n_ introduces an interesting feature of the *Rewired Model*.

Despite a poor match to experimental data and the *TM Model* with respect to DHF_n_, this proposed set of drug targets may replicate the clinical effect of TM and Sulfamethoxazole. Paradoxically, the drop in DHF_n_ replicates the task of the drug Sulfamethoxazole, which is used clinically with TM (the target of this drug target search effort). Sulfamethoxazole targets the enzyme DHFS, which synthesizes DHF_1_ and therefore inputs mass into the folate network. Sulfamethoxazole and TM clinically work together to fully shut down flux into and within the folate network [10, 16, 20]. The proposed inhibitors in the *Rewired Model* also appear to severely lower the amount of cellular DHF_n_, which is achieved clinically by Sulfamethoxazole. This means that the proposed set of inhibitors would be effective against some bacterial strains that show resistance to TM and Sulfamethoxazole simultaneously. This is an unexpected but welcome output of the *Rewired Model* which puts the poor SSE score in context and bolsters the result as a viable set of drug targets.

## Conclusions

The current work presents a mathematical model of the *E. coli* folate network that improves on the previous effort in accuracy, scope, and applicability. A detailed look at TM’s mechanism not only shows the importance of folate polyglutamation levels, but also of enzyme-metabolite interactions to overall dynamics of the folate cycle. The improved look at this well-studied system allows a programmatic drug target search for alternatives to both TM and Sulfamethoxazole. We present this approach as a radically more efficient method to dealing with antibiotic-resistant bacteria by building on past successes.

## Acknowledgements

We thank Rick Dilling and Vivian Callier for their valuable comments.

## Supporting information

S1 Table. TM Model parameters.

S2 Table. SSE Outputs (observations vs simulations).

S3 Table. Experimental Data Replicates.

S1 Fig. Network Rewiring Reaction Velocities.

General flux in the system is lower than in the TM Model, and resultant fluxes over time differ greatly from that of the TM Model.

## References

1. Andersson, D.I. and D. Hughes, Antibiotic resistance and its cost: is it possible to reverse resistance? Nat Rev Micro, 2010. 8(4): p. 260–271.

2. Stamm, L.V., Global Challenge of Antibiotic-Resistant Treponema pallidum. Antimicrobial Agents and Chemotherapy, 2010. 54(2): p. 583–589.

3. Gandhi, N.R., et al., Multidrug-resistant and extensively drug-resistant tuberculosis: a threat to global control of tuberculosis. The Lancet. 375(9728): p. 1830–1843.

4. El-Halfawy, O.M. and M.A. Valvano, Chemical Communication of Antibiotic Resistance by a Highly Resistant Subpopulation of Bacterial Cells. PLoS ONE, 2013. 8(7): p. e68874.

5. Baccanari, D.P., et al., Escherichia coli dihydrofolate reductase: isolation and characterization of two isozymes. Biochemistry, 1977. 16(16): p. 3566–3572.

6. Nijhout, H.F., et al., A mathematical model of the folate cycle: new insights into folate homeostasis. J Biol Chem, 2004. 279(53): p. 55008–16.

7. Roje, S., S-Adenosyl-l-methionine: Beyond the universal methyl group donor. Phytochemistry, 2006. 67(15): p. 1686–1698.

8. Miller, B.A. and E.B. Newman, Control of serine transhydroxymethylase synthesis in Escherichia coli K12. Canadian Journal of Microbiology, 1974. 20(1): p. 41–47.

9. Greene, R.C. and C. Radovich, Role of methionine in the regulation of serine hydroxymethyltransferase in Eschericia coli. Journal of Bacteriology, 1975. 124(1): p. 269–278.

10. Kwon, Y.K., et al., A domino effect in antifolate drug action in Escherichia coli. Nat Chem Biol, 2008. 4(10): p. 602–608.

11. Kwon, Y.K., M.B. Higgins, and J.D. Rabinowitz, Antifolate-induced depletion of intracellular glycine and purines inhibits thymineless death in E. coli. ACS Chem Biol, 2010. 5(8): p. 787–95.

12. Trimmer, E.E., et al., Folate activation and catalysis in methylenetetrahydrofolate reductase from Escherichia coli: roles for aspartate 120 and glutamate 28. Biochemistry, 2001. 40(21): p. 6216–26.

13. McGuire, J.J., et al., Enzymatic synthesis of folylpolyglutamates. Characterization of the reaction and its products. Journal of Biological Chemistry, 1980. 255(12): p. 5776–5788.

14. Shane, B., Pteroylpoly(gamma-glutamate) synthesis by Corynebacterium species. Purification and properties of folypoly(gamma-glutamate) synthetase. Journal of Biological Chemistry, 1980. 255(12): p. 5655–5662.

15. Lee, J.C., et al., The prevalence of trimethoprim-resistance-conferring dihydrofolate reductase genes in urinary isolates of Escherichia coli in Korea. Journal of Antimicrobial Chemotherapy, 2001. 47(5): p. 599–604.

16. Blahna, M.T., et al., The role of horizontal gene transfer in the spread of trimethoprim– sulfamethoxazole resistance among uropathogenic Escherichia coli in Europe and Canada. Journal of Antimicrobial Chemotherapy, 2006. 57(4): p. 666–672.

17. Aflaki, E., et al., Macrophage Models of Gaucher Disease for Evaluating Disease Pathogenesis and Candidate Drugs. Science Translational Medicine, 2014. 6(240): p. 240ra73.

18. Patnaik, S., et al., Discovery, structure-activity relationship, and biological evaluation of noninhibitory small molecule chaperones of glucocerebrosidase. J Med Chem, 2012. 55(12): p. 5734–48.

19. Bonhoeffer, S., M. Lipsitch, and B.R. Levin, Evaluating treatment protocols to prevent antibiotic resistance. Proceedings of the National Academy of Sciences, 1997. 94(22): p. 12106–12111.

20. Masters, P.A., et al., TRimethoprim-sulfamethoxazole revisited. Archives of Internal Medicine, 2003. 163(4): p. 402–410.

21. Gangjee, A., et al., Potent Dual Thymidylate Synthase and Dihydrofolate Reductase Inhibitors: Classical and Nonclassical 2-Amino-4-oxo-5-arylthio-substituted-6-methylthieno[2,3-d]pyrimidine Antifolates. Journal of medicinal chemistry, 2008. 51(18): p. 5789–5797.

22. Contestabile, R., et al., l-Threonine aldolase, serine hydroxymethyltransferase and fungal alanine racemase. A subgroup of strictly related enzymes specialized for different functions. Eur J Biochem, 2001. 268(24): p. 6508–25.

23. Kisliuk, R., Pteroylpolyglutamates. Molecular and Cellular Biochemistry, 1981. 39(1): p. 331–345.

24. Kisliuk, R.L., et al., Polyglutamyl derivatives of tetrahydrofolate as substrates for Lactobacillus casei thymidylate synthase. Biochemistry, 1981. 20(4): p. 929–934.

25. Horiuchi, Y., et al., Coupling effects of distal loops on structural stability and enzymatic activity of Escherichia coli dihydrofolate reductase revealed by deletion mutants. Biochimica et Biophysica Acta (BBA) - Proteins and Proteomics, 2010. 1804(4): p. 846–855.

26. Sheng, Y., et al., Mutation of an essential glutamate residue in folylpolyglutamate synthetase and activation of the enzyme by pteroate binding. Arch Biochem Biophys, 2002. 402(1): p. 94–103.

27. Bognar, A., et al., Folylpoly-gamma-glutamate synthetase-dihydrofolate synthetase. Cloning and high expression of the Escherichia coli folC gene and purification and properties of the gene product. J Biol Chem, 1985. 260: p. 5625 – 5630.

28. Bognar, A.L. and B. Shane, [55] Bacterial folypoly([gamma]-glutamate) synthase-dihydrofolate synthase, in Methods in Enzymology, D.B.M. Frank Chytil, Editor. 1986, Academic Press. p. 349–359.

29. Burton, E., J. Selhub, and W. Sakami, The substrate specificity of 5-methyltetrahydropteroyltriglutamate-homocysteine methyltransferase. Vol. 111. 1969. 793–795.

30. Whitfield, C.D., E.J. Steers, Jr., and H. Weissbach, Purification and properties of 5-methyltetrahydropteroyltriglutamate-homocysteine transmethylase. J Biol Chem, 1970. 245(2): p. 390–401.

31. Trimmer, E.E., et al., Aspartate 120 of Escherichia coli methylenetetrahydrofolate reductase: evidence for major roles in folate binding and catalysis and a minor role in flavin reactivity. Biochemistry, 2005. 44(18): p. 6809–22.

32. McGuire, J. and J. Bertino, Enzymatic synthesis and function of folylpolyglutamates. Molecular and Cellular Biochemistry, 1981. 38(1): p. 19–48.

33. Mansouri, A., J.B. Decter, and R. Silber, Studies on the regulation of one-carbon metabolism. II. Repression-derepression of serine hydroxymethyltransferase by methionine in Escherichia coli 113-3. J Biol Chem, 1972. 247(2): p. 348–52.

